# Distinct types of short open reading frames are translated in plant cells

**DOI:** 10.1101/213736

**Authors:** Igor Fesenko, Ilya Kirov, Andrey Kniazev, Regina Khazigaleeva, Vassili Lazarev, Daria Kharlampieva, Ekaterina Grafskaia, Viktor Zgoda, Ivan Butenko, Georgy Arapidi, Anna Mamaeva, Vadim Ivanov, Vadim Govorun

## Abstract

Genomes contain millions of short (<100 codons) open reading frames (sORFs), which are usually dismissed during gene annotation. Nevertheless, peptides encoded by such sORFs can play important biological roles, and their impact on cellular processes has long been underestimated. Here, we analyzed approximately 70,000 transcribed sORFs in the model plant *Physcomitrella patens* (moss). Several distinct classes of sORFs that differ in terms of their position on transcripts and the level of evolutionary conservation are present in the moss genome. Over 5000 sORFs were conserved in at least one of ten plant species examined. Mass spectrometry analysis of proteomic and peptidomic datasets suggested that 584 sORFs located on distinct parts of mRNAs and long non-coding RNAs (lncRNAs) are translated, including 73 conservative sORFs. Translational analysis of the sORFs and main ORFs at a single locus suggested the existence of genes that code for multiple proteins and peptides with tissue-specific expression. Alternative splicing is likely involved in the excision of translatable sORFs from such transcripts. We identified a group of sORFs homologous to known protein domains and suggested they function as small interfering peptides. Functional analysis of candidate lncRNA-encoded peptides showed it to be involved in regulating growth and differentiation in moss. The high evolutionary rate and wide translation of sORFs suggest that they may provide a reservoir of potentially active peptides and their importance as a raw material for gene evolution. Our results thus open new avenues for discovering novel, biologically active peptides in the plant kingdom.

## INTRODUCTION

The genomes of nearly all organisms contain hundreds of thousands of short open reading frames (sORFs; <100 codons) whose coding potential has been the subject of recent reviews (Andrews and Rothnagel 2014; Couso 2015; Hellens et al. 2016; Couso and Patraquim 2017). However, gene annotation algorithms are generally not suited for dealing with sORFs because short sequences are unable to obtain high conservation scores, which serve as an indicator of functionality (Ladoukakis et al. 2011). Nevertheless, using various bioinformatic approaches, sORFs with high coding potential have been identified in a range of organisms including fruit flies, mice, yeast and *Arabidopsis thaliana* (Ladoukakis et al. 2011; Hanada et al. 2013; Aspden et al. 2014; Bazzini et al. 2014). The first systematic study of sORFs was conducted on baker’s yeast, where 299 previously non-annotated sORFs were identified and tested in genetic experiments (Kastenmayer et al. 2006). Subsequently, 4561 conserved sORFs were identified in the genus *Drosophila*, 401 of which were postulated to be functional, taking into account their syntenic positions, low *K*_*A*_*/K*_*S*_ (<0.1) values and transcriptional evidence (Ladoukakis et al. 2011). In a recent study, Mackowiak and colleagues predicted the presence of 2002 novel conserved sORFs (from 9 to 101 codons) in *H. sapiens, M. musculus, D. rerio, D. melanogaster* and *C. elegans* (Mackowiak et al. 2015). The first comprehensive study of sORFs in plants postulated the existence of thousands of sORFs with high coding potential in Arabidopsis (Lease and Walker 2006; Hanada et al. 2007; Hanada et al. 2013), including 49 that induced various morphological changes and had visible phenotypic effects.

Recent studies have pointed to the important roles of sORF-encoded peptides (SEPs) in cells (Magny et al. 2013; Nelson et al. 2016; D’Lima et al. 2017; Huang et al. 2017; Matsumoto et al. 2017). However, unraveling the roles of SEPs is a challenging task, as is their detection at the biochemical level. In animals, SEPs are known play important roles in a diverse range of cellular processes (Kondo et al. 2010; Magny et al. 2013). By contrast, only a few functional SEPs have been reported in plants, including POLARIS (PLS; 36 amino acids), EARLY NODULIN GENE 40 (ENOD40; 12, 13, 24 or 27 amino acids), ROTUNDIFOLIA FOUR (ROT4; 53 amino acids), KISS OF DEATH (KOD; 25 amino acids), BRICK1 (BRK1; 84 amino acids), Zm-908p11 (97 amino acids) and Zm-401p10 (89 amino acids) (Andrews and Rothnagel 2014; Tavormina et al. 2015). These SEPs help modulate root growth and leaf vascular patterning (Chilley et al. 2006), symbiotic nodule development (Djordjevic et al. 2015), polar cell proliferation in lateral organs and leaf morphogenesis (Narita et al. 2004), and programmed cell death (apoptosis) (Blanvillain et al. 2011).

To date, functional sORFs have been found in a variety of transcripts, including untranslated regions of mRNA (5′ leader and 3′ trailer sequences), lncRNAs, and microRNA transcripts (pri-miRNAs) (Andrews and Rothnagel 2014; Laing et al. 2015; Lauressergues et al. 2015; Couso and Patraquim 2017). Evidence for the transcription of potentially functional sORFs has been obtained in *Populus deltoides, Phaseolus vulgaris, Medicago truncatula, Glycine max* and *Lotus japonicus* (Guillen et al. 2013). The transcription of sORFs can be regulated by stress conditions and depends on the developmental stage of the plant (De Coninck et al. 2013; Hanada et al. 2013; Rasheed et al. 2016). Indeed, sORFs might represent an important source of advanced traits required under stress conditions. During stress, genomes undergo widespread transcription to produce a diverse range of RNAs (Kim et al. 2010; Mazin et al. 2014); therefore, a large portion of sORFs becomes accessible to the translation machine for peptide production. Stress conditions can lead to the transcription of sORFs located in genomic regions that are usually non-coding (Giannakakis et al. 2015). Such sORFs appear to serve as raw materials for the birth and subsequent evolution of new protein-coding genes (Couso and Patraquim 2017).

The transcription of an sORF does not necessarily indicate that it fulfills any biological role, as opposed to being a component of the so-called translational noise (Guttman et al. 2013). According to ribosomal profiling data, thousands of lncRNAs display high ribosomal occupancy in regions containing sORFs in mammals (Ingolia et al. 2011; Aspden et al. 2014; Bazzini et al. 2014). However, lncRNAs can have the same ribosome profiling patterns as canonical non-coding RNAs (e.g., rRNA) that are known not to be translated, implying that these lncRNAs are unlikely to produce functional peptides (Guttman et al. 2013). In addition, identification of SEPs via mass spectrometry analyses has found many fewer peptides than predicted sORFs (Slavoff et al. 2013; Aspden et al. 2014). Thus, the abundance, lifetime and other features of SEPs are generally unclear.

We performed a comprehensive analysis of the sORFs that have canonical AUG start codons and high coding potential in the *Physcomitrella patens* genome. The translation of hundreds of sORFs was confirmed by mass-spectrometry analysis. From these, candidate lncRNA-encoded peptides were selected for further analysis, which provided evidence for their biological functions.

## RESULTS

### Discovery and classification of potential coding sORFs in the moss genome

Our approach is summarized in Fig. 1A. At the first stage of analysis, we used the sORFfinder tool (Hanada et al. 2010) to identify single-exon sORFs starting with an AUG start codon and less than 300 bp long. This approach resulted in the identification of 638,439 sORFs with high coding potential (CI index) in all regions of the *P. patens* genome.

**Fig. 1.**
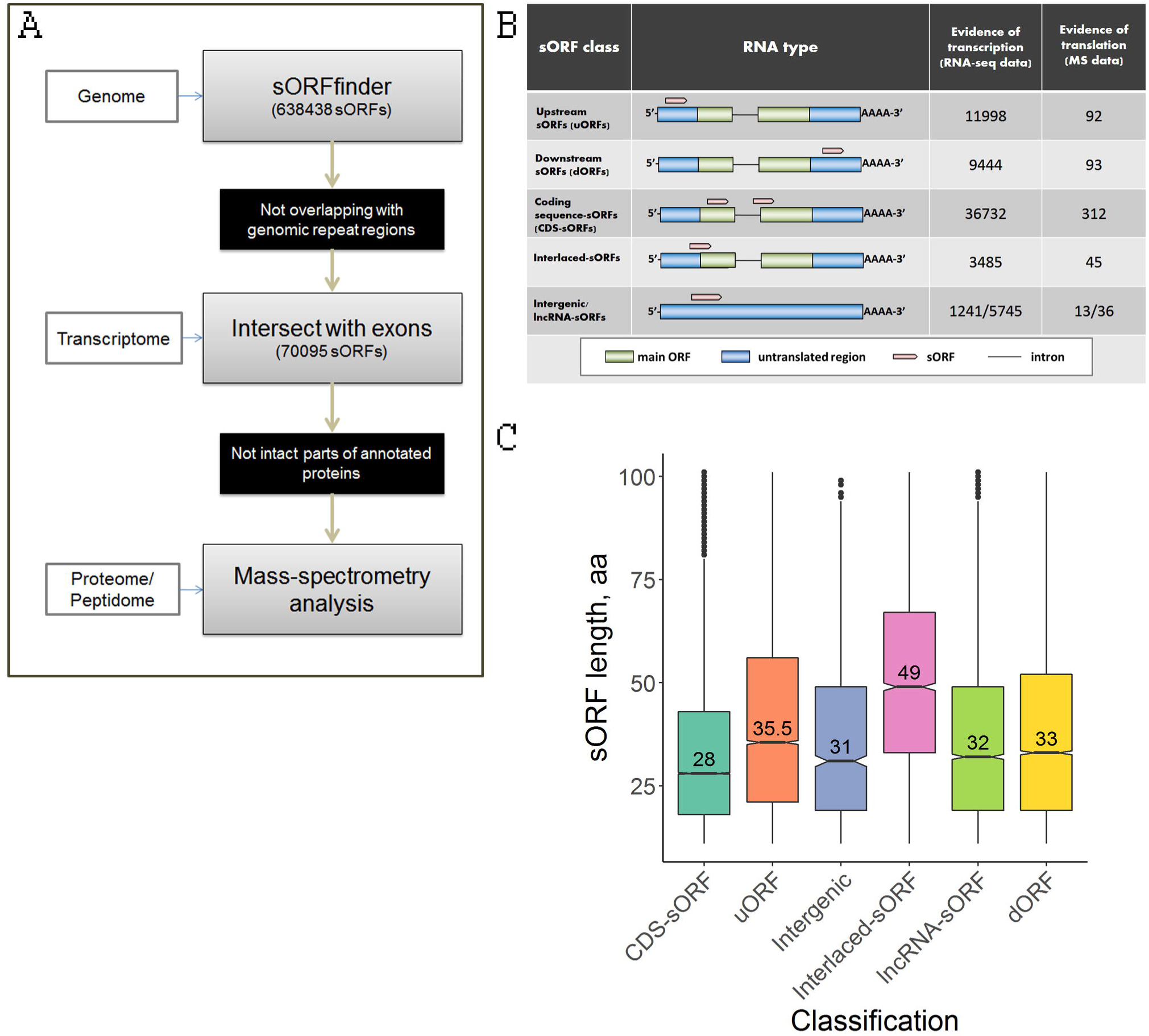
Several distinct types of sORFs are present in the moss genome. **A** – Pipeline used in this study to identify coding sORFs; **B** – Proposed classification of sORFs according to the types of encoding transcripts: upstream ORFs (uORFs) and downstream ORFs (dORFs) in the untranslated regions (UTRs) of canonical mRNAs; CDS-sORFs, which overlap with protein-coding sequences in alternative (+2 or +3) reading frames or are truncated versions of proteins generated by alternative splicing; interlaced-sORFs, which overlap both the protein-coding sequence and UTR on the same transcript; lncRNA-sORFs and intergenic sORFs, which are located on short non-protein coding transcripts.; **C** – Boxplot of the length distribution of sORFs in different groups; **D** – The results of GO enrichment analysis for genes possessing uORFs and CDS-sORFs. BP, CC and MF represent “Biological process”, “Cellular component” and “Molecular function”, respectively.

We selected 70,095 unique sORFs located on transcripts annotated in the moss genome (phytozome.jgi.doe.gov) and/or our dataset (Fesenko et al. 2015) for further analysis, as well as those on lncRNAs from two databases-CantataDB (Szczesniak et al. 2016) and GreenC (Paytuvi Gallart et al. 2016); sORFs located in repetitive regions were discarded (Supplemental Table S1). These selected sORFs, which were 33 to 303 bp long, were located on 33,981 transcripts (22,969 genes), with up to 28 sORFs per transcript (Supplemental Fig. S1A).

We then classified the sORFs based on their location on the transcript: 63,109 “genic-sORFs” (located on annotated transcripts, but not on lncRNA), 1241 “intergenic-sORFs” (located on transcripts from our dataset and not annotated in the current version of the genome) and 5745 “lncRNA-sORFs” (located on lncRNAs from CantataDB (Szczesniak et al. 2016), GreenC (Paytuvi Gallart et al. 2016) or our data set (Fesenko et al. 2017); Fig. 1B). The genic-sORFs include 11,998 upstream ORFs (uORFs; for 5’-UTR location), 9443 downstream ORFs (dORFs; for 3’-UTR location), 36,731 coding sequence-sORFs (CDS-sORFs; sORFs overlapping with main ORFs (+1 frame) in non-canonical +2 and +3 reading frames) and 3485 interlaced-sORFs (overlapping with both the CDS and 5’-UTR or CDS and 3’-UTR on the same transcript) (Fig. 1B, Supplemental Fig. S1B).

As expected based on the sORFfinder search strategy (Hanada et al. 2010), the sORF set was enriched in CDS-sORFs (52%, Fisher’s exact test, P-value = 1.736392e-285), whereas dORFs, uORFs and interlaced-sORFs were underrepresented (Fisher’s exact test, P-value < 4.792689e-88) compared to a random exonic fragments (REF) set, which was used as a negative control.

On average, CDS-sORFs (median size of 22 codons) were shorter than uORFs (median size of 35 codons; Mann-Whitney *U* test P = 2.2e-151) and dORFs (median length 32 codons, Mann-Whitney *U* test P = 1.03e-43). The median size of interlaced-sORFs was 49 codons, which is significantly longer than other genic-sORFs (Mann-Whitney *U* test P = 0.0021) (Fig. 1C).

Genes possessing CDS-sORFs were enriched in GO terms associated with protein binding and transferase activity, while genes possessing uORFs were enriched for signal transduction and transcriptional regulation (Supplemental Fig. S2). Such contrasting functional associations could be reflective of evolutionary trends that result in distinct sORF groups within protein coding genes.

### Analysis of evolutionary conservation of sORFs

It is widely accepted that evolutionary conservation is a strong indicator of functionality (Ladoukakis et al. 2011). To estimate the number of conserved sORFs in the moss genome and the evolutionary pressure on their amino acid sequences, we performed a tBLASTn search (e-value cutoff 0.00001) of each sORF sequence against the reconstructed genomes of three *P. patens* ecotypes: Villersexel, Reute, and Kaskasia as well as the transcriptomes of ten other species (Supplemental Fig. S3). The selected species include those that diverged from *P. patens* 177 (*Ceratodon purpureus*), 320 (*Sphagnum fallax*), 493 (*Marchantia polymorpha*), 532 (Arabidopsis thaliana, *Oryza sativa, Zea mays, Selaginella moellendorffii and Spirodela polyrhiza*) and 1160 (*Volvox carteri and Chlamydomonas reinhardtii*) Mya (According to www.timetree.org (Kumar et al. 2017); Supplemental Fig. S3).

A conservation analysis of the sORFs in the reconstructed genomes of these ecotypes showed that 2.4% (1618) of the sORFs were lacking either the start or stop codons in at least one species. Interestingly, CDS-sORFs (506) were significantly underrepresented in this set (Fisher’s exact test P-value < 2.2e-16), while uORF (478), dORF (278), lncRNA-sORFs (202), and intergenic-sORFs (59) were significantly overrepresented (Fisher’s exact test P-value < 2.2e-16). These results suggest that uORFs, dORF, lncRNA-sORFs, and intergenic-sORFs are prone to a shorter retention time than CDS-ORFs.

We found 5034 conserved sORFs with detectable homologous sequences in at least one species: 4797 in *C. purpureus*, 1049 in *S. fallax*, 436 in *M. polymorpha*, 328 in *S. moellendorffii*, 297 in *S. polyrhiza*, 275 in A. thaliana, 282 in *Z. mays*, 274 in *O. sativa*, 86 in *V. carteri* and 89 in *C. reinhardtii*. The number of conserved sORFs was negatively correlated with the time since divergence, with the fewest homologous sequences found in *V. carteri* and *C. reinhardtii*, which diverged more than 1000 Mya from a common ancestor. We found that lncRNA-sORFs were underrepresented among sORFs having homologs in the ten species examined (Fig. 2A). We also found significantly fewer uORFs and dORFs in the two closest species, *C. purpureus* and *S. fallax*, whereas CDS-sORFs were significantly overrepresented in these species (Fisher’s exact test, P<2.2e-16) (Fig. 2B).

**Fig. 2.**
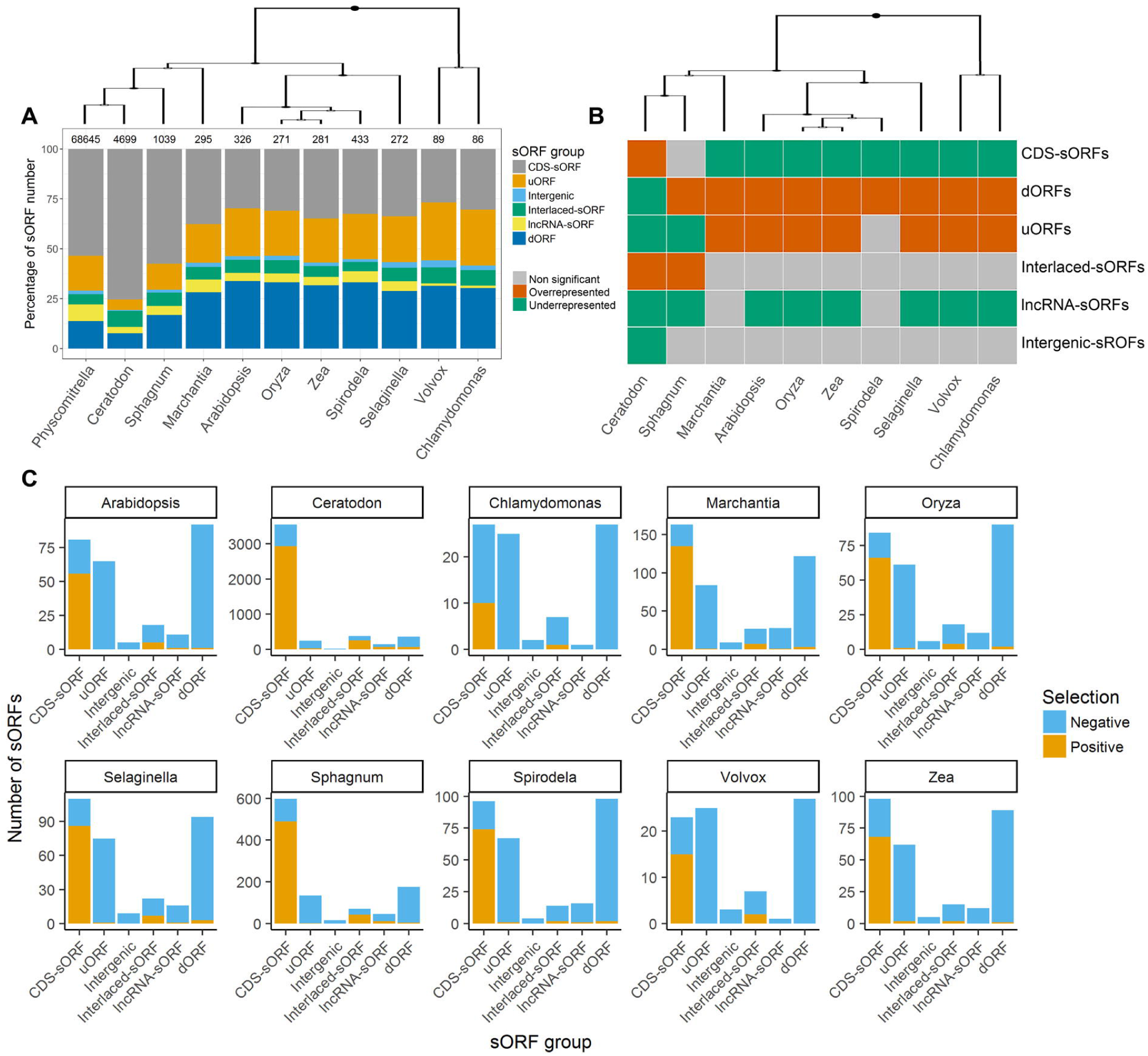
Analysis of the trends in sORFs evolution. A – The percentage of each type of sORF among sORFs having homologs in ten plant species. B – Statistical analysis (by Fisher’s exact test) of differences between the number of conservative sORFs in each of ten species and the initial dataset; C – Pairwise *K*_*A*_/*K*_*S*_ ratio distribution for each type of sORF conserved among ten plant species.

However, the portion of uORFs and dORFs found in the more distant species was increased relative to the initial dataset compared to CDS-sORFs, causing their significant overrepresentation (Fisher’s exact test, P<0.0005). Thus, the relative enrichment of conserved CDS-sORFs and interlaced-sORFs found in the two closest species of *P. patens, C. purpureus* and *S. fallax*, resulted from a significant reduction in the number of uORFs and dORFs (Fig. 2A). As a control, we also investigated changes in the proportion of uREFs, dREFs and CDS-REFs in these ten species and obtained opposite results (Supplemental Fig. S4). To compare this trend with that of the protein coding genes, we selected 158 annotated *P. patens* genes that code for small proteins without introns (< 100 aa). The percentages of sORFs and these proteins showing homology with at least one species were significantly different (7.2% sORFs vs. 86% small proteins), pointing to high genome turnover of sORF sequences. We next assessed whether the lengths of homologous sORFs from other species were the same as those in moss or if they varied in size. According to our data, most putative homologous sORFs differed in length, contributing to sORF diversification (Supplemental Fig. S5).

To better understand the large-scale trends of sORF evolution, we examined the differences in selection pressure at the amino acid level between different major groups of sORFs (CDS-sORFs, uORFs, dORFs, lncRNA-sORFs, interlaced-sORFs) using the criterion of *K*_*A*_/*K*_*S*_. This analysis showed that the highest portion of sORFs comprised CDS-sORFs, with *K*_*A*_/*K*_*S*_ ratio > 1, implying ongoing positive selection of sORFs emerging in the CDS of protein-coding genes. This criterion for other sORF groups was < 1 in most cases, pointing to purifying selection for these sequences (Fig. 2C).

Thus, evolutionary analysis demonstrated that the conservation of an sORF on a large evolutionary scale differs from that of randomly selected exon sequences and depends on the location of the sORF. Higher retention rates were observed for uORFs and dORFs, whereas CDS-sORFs and lncRNA-ORFs were under strong positive selection.

### Experimental evidence for the translation of sORFs

Obtaining evidence for the translation of sORFs is an important step towards identifying functional SEPs. We analyzed the Kozak consensus sequences (Kozak 1986) surrounding sORF start codons. Kozak consensus sequence plays an important role in translation initiation (Kozak 1997). Depending on the presence of the purine in position −3 and the G in position +4 (where +1 is “A” in the “AUG” codon) the Kozak was considered to be “strong” (both are present), “medium” (one is present) or “weak” (neither are present) (Kozak 1997). According to our results, 41816 (∼60%) of the predicted sORFs were surrounded by “strong” and “medium” Kozak sequences. These values were significantly smaller than those of annotated protein coding ORFs (87%, Fisher’s exact test P-value < 2.2e-16).

We then verified the translation of our predicted sORFs using mass-spectrometry (MS) analysis. Taking into account the shortage of proteomic methods for identifying small proteins or peptides, in the current study, we generated two datasets: the “peptidomic” dataset - endogenous peptides extracted from three types of moss cells: gametophores, protonemata and protoplasts and the “proteomic” dataset - tryptic peptides generated in a standard proteomic pipeline (Supplemental Table S2). All datasets were mapped with MaxQuant against a custom database containing our sORFs together with nuclear, chloroplast and mitochondrial moss protein sequences (see details in the Methods). PSMs (peptide spectrum matches) were identified at 1 % FDR, and ambiguous peptides were filtered out. In total, we confirmed the translation of 584 sORFs: 198 in gametophores, 277 in protonemata, and 190 in protoplasts (Fig. 3A, Supplemental Table S3). These results indicate tissue-specific translation of sORFs. The most prominent group of translatable sORFs consisted of CDS-sORFs (305, 51%) (Fig. 3B). Interestingly, the translation of 36 sORFs located on lncRNAs was also detected by our analysis. Approximately 60% of the translated sORFs (349 sORFs) contained “strong” and “medium” Kozak elements, which is a similar to the results obtained for all predicted sORFs (∼60%). This result suggests that translation initiation may differ for sORFs and protein coding ORFs.

**Fig. 3.**
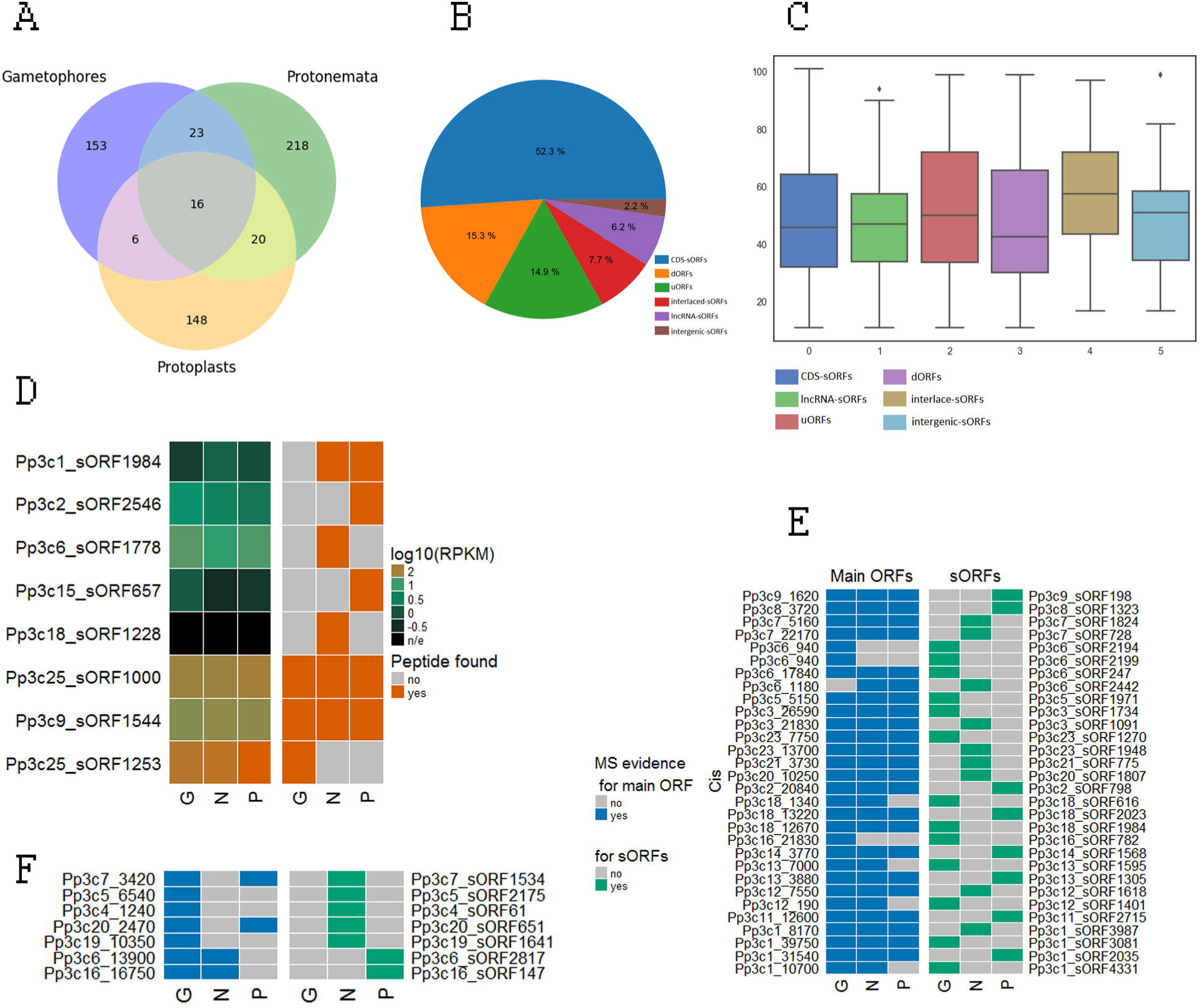
Moss contains hundreds of translatable sORFs. A – Venn diagram showing the distribution of the identified sORFs among three types of moss cells; B - Distribution of translatable sORFs based on the suggested classification; C - Length distribution of various groups of translatable sORFs; D - Heatmap showing expression levels (log10(RPKM)) for the lncRNAs (left) carrying sORFs (lncRNA-sORFs) and binary heatmap showing evidence of translation (determined as whether a peptide was identified (brown) or not (grey) in MS data) for the corresponding lncRNA-sORFs (right) in three moss tissues, gametophores (G), protonemata (N) and protoplasts (P); E – Binary heatmap showing evidence of translation for sORFs and proteins in multicoding genes in three moss tissues. G, N and P correspond to gametophores, protonemata and protoplasts, respectively. F - Examples of contrasting translational patterns of the main ORF and CDS-sORF. Only proteins confirmed by more than three unique tryptic peptides in the MS data are shown.

The length of translatable sORFs ranged from 11 to 100 amino acids (aa), which were generally longer than untranslatable sORFs (Mann-Whitney *U* test P = 4e-53) (Fig. 3C). The length of interlaced-sORFs differed significantly from that of CDS-sORFs and lncRNA-sORFs (Mann-Whitney *U* test P = 0.002 and Mann-Whitney *U* test P = 0.001, respectively) but did not differ from uORFs (Mann-Whitney *U* test P = 0.06). We observed that PSMs supporting SEP identifications had lower average quality than those mapped to the protein sequences of all datasets (Supplemental Figs. S6A and S6B). This finding is in agreement with data obtained for the animal kingdom (Slavoff et al. 2013; Mackowiak et al. 2015). The quality of spectra and the values of PSMs supporting the expression of SEPs were better in the “peptidomic” dataset (Supplemental Fig. S6C). Also, translatable sORFs were longer for those identified in the peptidomic dataset (Supplemental Fig. S6D).

There were no significant dependencies between the level of expression of a transcript and the chance of finding peptides from sORFs located on this transcript (logistic regression, P-value >> 0.05). However, among the 16 sORFs with evidence of translation in all types of moss cells, lncRNA-sORFs were significantly overrepresented (Fisher’s exact test, P-value = 0.001). Two of these SEPs, Pp3c9_sORF1544 (41aa) and Pp3c25_sORF1000 (61aa), were common to all three cell types and were confirmed by 15 and 17 unique endogenous peptides, respectively (Fig. 3D). The level of transcription of some lncRNAs (according to the previous data (Fesenko et al. 2017) and evidences of translation for the corresponding lncRNA-sORFs are shown in Fig. 3D. These data may point to biological significance for the peptides translated from these sORFs rather than the sORFs having regulatory functions in the translation of the main ORF. To explore this notion, we investigated the functions of three SEPs encoded by lncRNAs (see below).

### sORFs can be translated together with proteins

Several reports provide evidence that eukaryotic mRNA can have more than one coding ORF (bi- and polycistronic genes) in both plants and animals (Blumenthal 1998; Rohrig et al. 2002; Pi et al. 2009; Tautz 2009; Xu et al. 2010). Based on our MS data, we identified 144 loci with at least two translated ORFs (annotated as main ORF and sORF), including 82 CDS-sORFs, that represent putative multi-coding genes (Supplemental Table S4). The translation of multiple ORFs can occur from either different transcripts of the same gene or consecutively from the single transcript (polycistronic transcript). Some of the putative multi-coding genes were translated simultaneously with protein-coding ORFs in the same type of moss cell (Fig. 3E), while others showed patterns of sORF and main ORF translation such that their products were present in different types of cells (Fig. 3F). This observation suggests that specific regulatory mechanisms may exist to fine-tune the translation of both sORFs and proteins situated in the same gene locus. Taken together, our findings indicate that at least 27% of translatable CDS-sORFs are expressed simultaneously with main ORFs and the translation of sORFs and proteins located together in the same locus might be regulated in a tissue-specific manner.

### Most translatable sORFs are not evolutionarily conserved

Analysis of the evolutionary conservation of sORFs is often a key step in revealing biologically active sORFs (Andrews and Rothnagel 2014). To determine whether the translatable sORFs were more highly conserved than the other sORFs, we analyzed the intactness of these sORFs in the reconstructed genomes of three *P. patens* ecotypes, ‘Villersexel’, ‘Reute’ and ‘Kaskasia’, as well as the ten abovementioned species. We found that 19 (3.3%) of 584 translatable sORFs in the ecotypes either lost the start/stop codon or had a frameshift or premature termination codon (PTC). This number was not significantly different from the number (2.4%, 1598 sORFs) for which translation was not detected by MS data, suggesting that sORF translation does not disrupt trends of sORF elimination in these ecotypes.

To investigate whether the trend in translatable sORF evolution differs from that of the other sORFs, we estimated the number of species in which homologs can be found and the selection pressure (*K*_*A*_/*K*_*S*_) on translatable sORFs on an evolutionary timescale using the transcriptomes of the ten abovementioned species. Overall, we found 73 sORFs had evidence of translation and conservation in at least one species while only 11 were under negative selection (*K*_*A*_/*K*_*S*_ ≪ 1) (Supplemental Fig. S7). Sixty-four (88%) of these were CDS-sORFs or interlaced-sORFs. These results suggest that these types of sORFs are more conserved. Although conservative sORFs were significantly enriched in a set of translatable sORFs (Fisher’s exact test, P = 2.716567e-05), we found that most translatable sORFs (87.6%) were not conserved.

We next examined whether the translatable sORFs detected in this study share similarity with a recently defined set of 13,748 putative SEPs in the *A. thaliana* (Hazarika et al. 2017). We identified two sORFs (Pp3c20_sORF627 (CDS-sORF), Pp3c11_sORF854 (CDS-sORF)) with evidence of translation according to our MS analysis that shared similarity with ARA-PEP peptides (e-value < 0.01), implying that these sORFs are evolutionarily conserved and may produce peptides in *A. thaliana* cells.

### Alternative splicing regulates the number of sORFs in protein-coding transcripts

Alternative splicing (AS) events may lead to the specific gain, loss or truncation of different groups of sORFs located on the transcripts of the same gene. For example, AS can generate sORFs that are truncated version of proteins (see below). We found 6092 alternatively spliced sORFs (AS-sORFs) belonging to transcripts from 4389 genes. CDS-sORFs were significantly overrepresented (Fig. 4A), while interlaced-sORFs, uORFs and dORFs were significantly underrepresented among AS-sORFs compared to the control dataset (AS-REF). The number of translatable sORFs in the set of AS-sORFs did not significantly differ from that expected by chance (Fisher’s exact test p-value=0.9423), suggesting that AS does not preferentially occur in peptide-encoding sORFs. Ten GO terms linked with nucleic acid binding (GO:0001071, GO:0003700), signal transducer activity (GO:0004871), aminopeptidase activity (GO:0004177), transferase activity (GO:0003950, GO:0016772, GO:0016775) and kinase activity (GO:0004672, GO:0004673, GO:0000155) were specifically enriched in a set of AS-sORF-carrying genes. These results demonstrate that AS-sORFs are located in regulatory genes more frequently than is expected by chance suggesting a potential role for sORFs in the translational regulation of these genes.

**Fig. 4.**
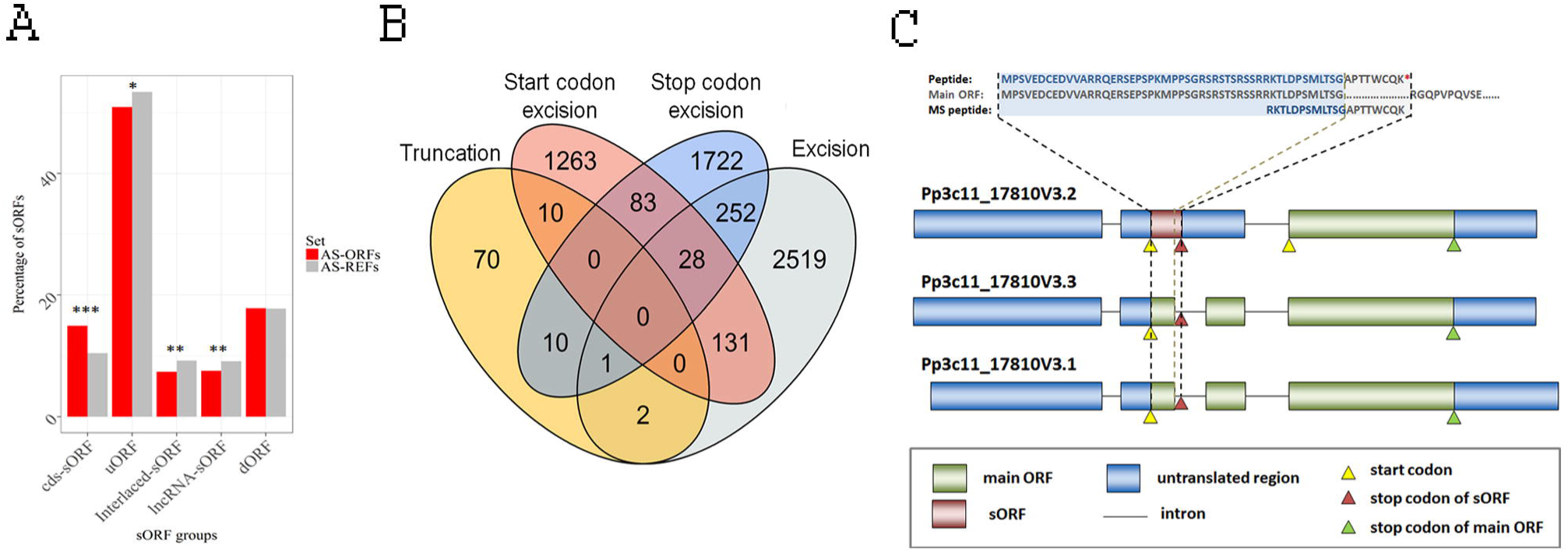
Alternative splicing regulates the expression of sORFs. A – Enrichment analysis of different sORF groups in a set of AS-sORFs and AS-REFs. P-value was calculated by Fisher’s exact test. ***P < 1e-10; **P < 0.001; *P < 0.05; B – Venn diagram showing the number of AS-sORFs influenced by different AS events. C – Example of a translatable CDS-sORF, which was generated by an AS event and partially overlaps with the main ORF of Pp3c11_17810. Intron retention caused the formation of the isoform with the sORF, while splicing of this intron led to the excision of the sORF stop codon and its disruption. MS detection of the peptide located at the exon-AS-intron junction allowed the translation of the sORF to be unambiguously distinguished from the translation of the main ORF. Upper panel shows the amino acid sequence of the sORF-encoded peptide, MS detected peptide and (partial) protein translated from the main ORF. Black and gray dotted lines mark the borders of the sORF and the canonical intron start site, respectively. The intron-exon structure of three transcript isoforms of the gene was retrieved from Phytozome (v12).

We randomly selected ten different translatable AS-sORFs and searched for the corresponding isoforms with/without sORFs in the transcriptomes of three types of moss cells. RT-PCR analysis revealed the transcription of these isoforms, confirming that they could indeed be translated (Supplemental Fig. S8). Moreover, four sORFs contained isoforms showing tissue-specific transcription. These observations led to the hypothesis that the translation of sORFs is regulated by AS.

We then classified the splicing events that lead to changes in sORF sequences into four groups: 1) truncation, when the middle region of the sORF was excised by splicing; 2) stop codon excision, when the sORF stop codon was spliced out; 3) start codon excision, when the sORF start codon was spliced out; and 4) excision, if the complete sORF was removed from an isoform. We found that half of the sORFs (48%, 2933) had undergone complete excision from their transcripts, whereas only 93 sORFs were truncated (1.5%) and 517 sORFs (12%) were affected by two or more events (Fig. 4B). Moreover, the complete excision of sORFs occurred significantly more frequently in uORFs than in the other sORF groups (57% vs. 20–44%, Fisher’s exact test P-value < 1e-05). In addition, evolutionarily conserved sORFs (conserved in >1 species) were significantly underrepresented in the set of AS-sORFs that were subject to complete excision (Fisher’s exact test P-value = 6.76e-42) compared to the other sets of AS-sORFs (“truncation”, “stop codon excision”, and “start codon excision”). Thus, our analysis demonstrated that AS leads to the excision of sORFs from the transcriptome of *P. patens*, preventing AS-sORF translation.

### The role of sORFs in modulating protein–protein interactions

Protein–protein interactions (PPI) are critical for the formation of higher order protein complexes. Competitive inhibitors of PPI are referred to as MicroProteins (miPs) or small interfering peptides (siPEPs) (Seo et al. 2011; Eguen et al. 2015). These proteins, which are usually small, can be generated by alternative splicing or evolutionarily generated by domain loss (Staudt and Wenkel 2011; Eguen et al. 2015). We hypothesized that sORFs with similarity to known proteins may compete with such proteins to impair their functions. To identify such sORFs, we performed BLASTP (E-value < e-5) similarity searches between the encoded amino acid sequences of sORFs and the annotated proteins of *P. patens*. We identified 363 sORFs resulting from AS events that partially overlapped with the main ORF, thereby generating truncated versions of the proteins. Based on the analogy of cis-miPs generated by alternative splicing events (Eguen et al. 2015), we will refer to these SEPs as cis-SEPs (and accordingly, cis-sORFs; Supplemental Table S5).

We analyzed how many cis-ORFs contained known complete or incomplete protein domains, finding that 60 sORFs harbored intrinsically disordered regions (IDRs, (van der Lee et al. 2014)), while 30 cis-sORFs contained parts of 28 different domains (Supplemental Table S5). The genes containing cis-sORFs were enriched in kinase and kinase-like domains. Among these, we observed the protein kinase domain (PS50011, Pp3c13_sORF653), protein tyrosine kinase (PF07714, Pp3c11_sORF2084) and MYB-like DNA-binding domain (TIGR01557, Pp3c19_sORF797). GO enrichment analysis also revealed significant overrepresentation of terms associated with protein modifications, such as GO:0006468 (protein phosphorylation) and GO:0036211 (protein modification process).

Among genes containing cis-sORFs, we identified some with similarity to putative transcription factor genes (TFs) such as genes encoding GROWTH-REGULATING FACTOR (e.g., Pp3c20_10590), C2H2 zinc finger domain containing (e.g., Pp3c1_16920), BTB/POZ domain containing (e.g., Pp3c16_9230), B3 DNA binding domain containing (e.g., Pp3c7_7990) and MYB-CC type transcription factor (e.g., Pp3c21_2850). Due to their similarity with TF domains, we predict they may act as dominant-negative repressors of TFs.

To obtain evidence for the translation of these sORFs, we analyzed MS data and found at least two examples (Fig. 4C). A few detected translatable cis-sORFs could be explained by a significant overlap with the protein sequences, whereas we filtered out the ‘ambiguous’ PSMs. Moreover, the formation of a premature termination codon (PTC) as a result of intron retention events, might lead to mRNA decay (Ge and Porse 2014; Karousis et al. 2016) and rapid nonsense-mediated decay (NMD)-coupled degradation of sORF-encoded peptides (Popp and Maquat 2013).

We identified 272 sORFs that shared similarity with annotated proteins but were located on other transcripts (trans-sORFs, see in Supplemental Table S5). The translation of six trans-sORFs was confirmed by our MS data. We found 36 potential trans-SEPs with similarity to known protein domains (Supplemental Table S5). Trans-sORFs may have originated through the divergence of ancient paralogous genes, which occurred after the paleo duplication of the moss genome (Rensing et al. 2007; Rensing et al. 2008). In fact, 159 (58.5%) trans-sORFs shared similarity to genes from at least one species. In addition, all of these trans-sORFs are under strong purifying selection (*K*_*A*_/*K*_*S*_ ≪ 1).

We then investigated which trans-sORFs share similarity to large gene families. Several distinct clusters with sORF-encoded peptides sharing similarity with more than four proteins from distinct genes were detected (Supplemental Fig. S9). Each cluster encompasses genes from different protein families, including one containing leucine-rich repeat and zinc-finger domains involved in protein–protein and protein–nucleic acid interactions, respectively.

To compete with target proteins, we presume that potential SEPs and their targets should coexist in a cell. We examined the co-expression data and compared the distribution of correlation coefficient values with those from randomly selected pairs (10 iterations) of genes. On average, these sORF-protein pairs had higher correlation coefficients than randomly selected gene pairs (Wilcoxon Rank Sum and Kolmogorov-Smirnov Tests P-value < 0.05), implying that sORF-bearing and target genes are frequently co-expressed.

### SEPs regulate moss growth

Despite the recent finding that 10% of overexpressed intergenic sORFs have clear phenotypes in Arabidopsis (Hanada et al. 2013), the functions of most sORFs and SEPs in plants are generally unknown. Known bioactive SEPs in plants are encoded by sORFs located on short non-protein-coding transcripts, which can be referred to as lncRNAs (Rohrig et al. 2002; Chilley et al. 2006). In this context, it would be intriguing to determine how many plant lncRNAs encode peptides, as well as the biological functions of these SEPs. Our pipeline allowed us to identify hundreds of translated sORFs, including those encoded by lncRNAs. Some of these lncRNA-sORFs showed tissue-specific transcription and translation patterns, while others were expressed in all types of moss cells (Fig. 3C). We reasoned that stably expressed lncRNA-sORFs can produce peptides that play fundamental roles in various cellular processes. To explore this hypothesis, we examined the impact of lncRNA-sORF overexpression and knockout on moss morphology using three conserved lncRNAs-sORFs: Pp3c9_sORF1544, Pp3c25_sORF1253, Pp3c25_sORF1000 (Fig. 3C). We obtained multiple independent mutant lines for each of these lncRNAs-sORFs (Supplemental Figs. S10 and S11). Both the overexpression and knockout of sORFs resulted in morphological changes, implying that these peptides play a role in growth and development of *P. patens* (Fig. 5).

**Fig. 5.**
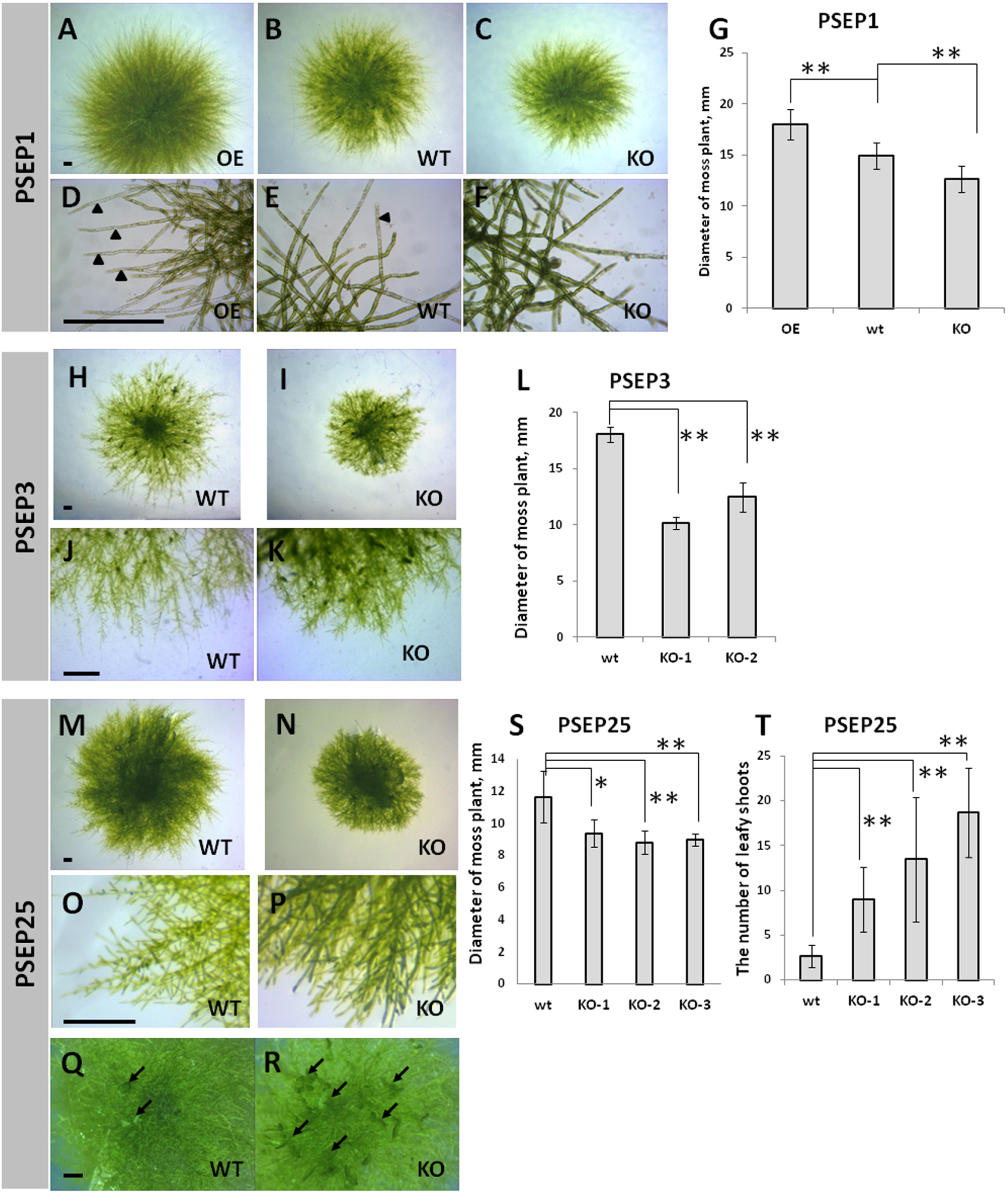
Morphology of wild type and sORF-encoded peptide mutant lines. **The phenotypes of PSEP1 knockout (KO) and overexpression (OE) lines grown on BCD medium with 0.5% glucose:** A, D – overexpression of PSEP1; C, F – knockout of PSEP1; G – the diameter of moss plants with overexpression and knockout of sORF-encoded peptide PSEP1. **The phenotypes of PSEP3 knockout (KO) lines grown on BCD medium:** H, J – wild type; I, K – knockout lines; L - the diameter of moss plants with knockout of PSEP3. **The phenotypes of PSEP25 knockout (KO) lines grown on BCDAT medium:** M, O, Q – wild type; N, P, R – knockout lines; S - the diameter of moss plants with knockout of PSEP25. T – the number of leafy gametophores in wild type and three PSEP25 knockout lines. Arrows show young leafy gametophores. Scale bar: 500 mm. P-value was calculated by Student’s t-test. **P < 0.01; *P < 0.05.

Overexpression of a 41-aa peptide (*PSEP1, Physcomitrella patens* sORF encoded peptide 1) encoded by the lncRNA-sORF Pp3c9_sORF1544 resulted in longer caulonema cells (filaments implicated in a rapid radial extension of the protonemal tissues) compared to the wild-type and knockout lines (Figs 5A-G, Supplemental Fig. S12). Moreover, there was a significant difference in growth rate between the wild-type and *psep1* mutant lines (Fig. 5G). Rapid growth in the *PSEP1* overexpressing lines was accompanied by earlier aging and cell death (Supplemental Fig. S13).

The lines with a knockout in a 57-aa peptide (*PSEP3*), encoded by lncRNA-sORF Pp3c25_sORF1253 displayed a decrease in growth rate, altered filament branching, and shorter lateral filaments compared to the wild type (Figs. 5H-L, Supplemental Fig. S12). Similar to the results for the *PSEP3* knockout, knocking out a 61-aa peptide (*PSEP25*) encoded by lncRNA-sORF Pp3c25_sORF1000 also resulted in a decreased in growth rate and altered protonemal architecture on cultural medium without glucose (Figs. 50-T). *PSEP25* knockouts also had an increase in the number of leafy shoots (Figs. 5Q, R and T).

Taken together, our findings suggest that lncRNA-sORFs can influence growth and development in moss.

## DISCUSSION

Although functionally characterized SEPs have been shown to play fundamental roles in key physiological processes, sORFs are arbitrarily excluded during genome annotation. Given the difficulty in identifying translatable, functional sORFs, we know little about their origin, evolution and regulation in the genome. In the present study, we investigated the abundance, evolutionary history and possible functions of sORFs in the genome of the model moss *Physcomitrella patens.* The use of an integrated pipeline that includes transcriptomics, proteomics, and peptidomics data allowed us to identify hundreds of translatable sORFs in three types of moss cells. We propose that several distinct classes of sORFs that differ in terms of their position on transcripts, the level of evolutionary conservation, and possible functions are present in the moss genome (Fig. 6).

**Fig. 6.**
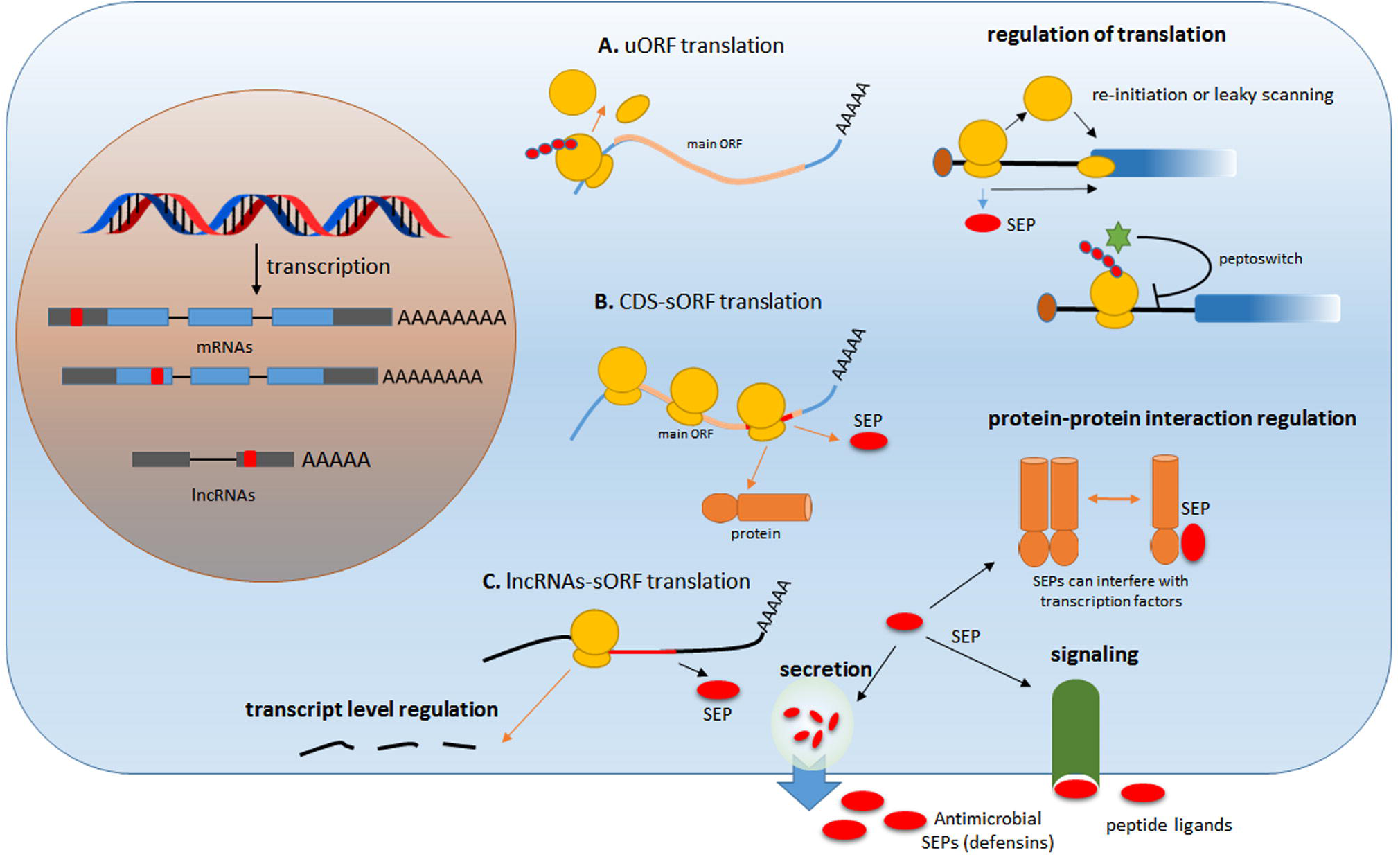
Proposed functions of sORF-encoded peptides. A – uORFs can function in the regulation of translation of the downstream main ORF. The functions of peptides encoded by uORFs are unknown, and most are likely to represent “noise” from protein translation; B – CDS-sORF-encoded peptides can help regulate protein-protein interactions, and some interfere with the translation of the main ORF; C – long non-coding RNAs or intergenic-sORFs can produce biologically active peptides that perform signaling, defense or regulatory functions. In addition, the translation of sORFs can activate the nonsense-mediated decay (NMD) mechanism, which leads to the degradation of the corresponding transcripts.

### sORFs with high coding potential are not conserved among genomes

Even though the analysis of sequence conservation is somewhat biased against the detection of short sequences (Ladoukakis et al. 2011), this technique is widely used to select candidate functional sORFs. Although analyzing the conservation of short amino acid sequences is not trivial (Moyers and Zhang 2016), hundreds of conserved sORFs have recently been identified in plants, yeast and animals (Ladoukakis et al. 2011; Hanada et al. 2013; Mackowiak et al. 2015). The number of sORFs conserved in the plant kingdom is undoubtedly underestimated due to the low sensitivity of tools used for conservation analysis and the limited number of available sequenced genomes from closely related species. Our pipeline allowed us to identify 5034 conserved sORFs among the transcriptomes of ten different plant species, 71 of which showed evidence of translation according to our MS data. However, we suggest that the possibly functional sORFs might significantly outnumber the conserved ones.

Despite the evidence for translation of approximately 1% of uORFs and dORFs, these sORF types were significantly underrepresented among the sORFs that are conserved in the closest related species. We even detected rapid inactivation of uORFs and dORFs in the reconstructed genomes of three *P. patens* ecotypes due to disruptions in the start or stop codons (47% of the total disrupted sORFs). As the occurrence of sORFs downstream or upstream of the main ORF can be deleterious to its translation, we cannot rule out the possibility that this may cause strong selection pressure and the rapid elimination of uORFs and dORFs (Iacono et al. 2005; Neafsey and Galagan 2007; Johnstone et al. 2016). Moreover, we observed significant depletion (Fisher’s exact test P-value = 5.25e-13) of uORFs and dORFs in a set of translatable conservative sORFs. Taken together, these findings suggest that sORFs located in untranslated regions are evolving rapidly and may play regulatory roles rather than encoding bioactive peptides.

In recent studies, thousands of alternative proteins were experimentally detected in human cell lines (Vanderperre et al. 2013; Samandi et al. 2017). In *P. patens*, we found tens of thousands of sORFs (CDS-sORFs) that overlapped with the CDS of protein-coding genes, 305 of which were translatable. The evolution of CDS-sORFs is undoubtedly an expensive process for the cell, as these elements may be located in regions encoding protein domains and influence the structure and function of the protein encoded by the main ORF (Cherry 2010). We found both CDS-sORFs originated from regions associated with known protein domains and CDS-sORFs from disordered regions, with higher conservation for CDS-sORFs originated from protein domain-encoding regions. These results indicate that the evolution of CDS-sORFs depends on their locations insight main CDS sequence.

In the current study, we found that both the transcription and translation of CDS-sORFs occurred in a tissue-specific manner. Protein-coding genes with tissue-specific transcription patterns and functional redundancy of the gene product are often under positive selection (Zhang and Li 2004; Montoya-Burgos 2011). This finding, together with other properties of CDS-sORFs, such as their overlap with particular parts of protein-coding sequences, might explain the high turnover rate of CDS-sORFs. However, whether sORFs are preferentially generated in fast-evolving regions of proteins or whether the selective pressure on sORFs leads to changes in protein-coding sequences is still unknown.

### Analysis of sORF translation: approaches that makes sense

It was recently suggested that sORFs are randomly generated in a genome (Couso and Patraquim 2017). Assuming that the average length of an sORF is approximately 60 bp and that sORFs do not overlap, these elements occupy a substantial portion of the moss genome. This raises the question: to what extent are sORFs present in the transcriptome and the proteome of a cell? According to ribosome profiling data from a wide variety of species, sORFs translation appears to occur in a pervasive manner (Ingolia et al. 2011; Guttman et al. 2013; Bazzini et al. 2014; Couso and Patraquim 2017). However, ribosome-profiling data alone are not sufficient to classify transcripts as coding or noncoding (Guttman et al. 2013). Thus, alternative approaches such as proteomics and peptidomics should be used to investigate the translation of sORFs (Slavoff et al. 2013; Ma et al. 2016). Mass-spectrometry studies have thus far confirmed the presence of a few dozen SEPs in the peptidomes of animal cells (Slavoff et al. 2013; Prabakaran et al. 2014; Mackowiak et al. 2015; Ma et al. 2016). Comparisons of ribosome profiling and mass spectrometry results have led to the conclusion that MS detects peptides arising from the most highly translated sORFs (Aspden et al. 2014; Bazzini et al. 2014). However, a recent study showed that there are no technical obstacles to the detection of sORF-encoded peptides by mass spectrometry (Verheggen et al. 2017).

In previous studies, only standard proteomics analysis was used to identify SEPs. We reasoned that analyzing endogenous peptide pools instead of tryptic peptides has several disadvantages in terms of SEP identification: 1) standard proteomic approaches are not suitable for the isolation and analysis of small and low-abundance peptide molecules; and 2) SEPs are shorter than standard proteins and it is unlikely that more than one tryptic fragment will be detected in a single proteomic experiment. Moreover, peptidomic approaches can theoretically be used to identify full-length SEPs in a cell. We firstly used endogenous peptides pools to detect SEPs and according to our data the values of PSMs, supporting expression of SEPs, were better in “peptidomic” dataset. Moreover, some SEPs were confirmed by several endogenous peptides (up to 17), that an increase the reliability of their detection. Notably, we did not observe any significant overlap between the sORFs detected using proteomic and peptidomic approaches. Thus, our study demonstrates the advantage of using complementary approaches for building a complete list of SEPs.

According to our MS data, the translation patterns of most sORFs tend to be tissue specific (Fig. 3A). We suggest that the slight overlap in tissue-specific expression among SEPs from various types of moss cells could be due to either specific SEP post-translational modification (PTM) patterns, tissue-specific transcription of sORFs, or the limitations of mass-spectrometry in detecting low-abundance or modified sORF-encoded peptides. According to our results, alternative splicing is an additional mechanism that control tissue-specific sORF expression in plant cells. Also, the number of sORFs that were commonly translated between two types of moss cells was higher for related cell types: protonemata and gametophores (two growth stages) as well as protonemata and protoplasts (protoplasts were generated from the protonemata). These observations indicate tissue-specific characteristics of SEPs translation and modification rather than technical limitation in detection.

### Functionality of SEPs

We identified hundreds of translatable sORFs representing multiple sORF types and suggested various functions for the types of sORFs (Fig. 6). Clear evidence of transcription and translation points to a possible biological significance of the sORFs that we identified here. Based on our conservation analysis and MS data, we suggest that the majority of uORFs and dORFs play regulatory roles instead of encoding peptides (Fig. 6A). By contrast, CDS-, interlaced- and lncRNA-sORFs have greater potential to encode bioactive peptides, as they are more highly conserved, frequently contain known protein domains and, according to the MS data, often produce peptides. However, the functions of these peptides are unclear and require more detailed investigation.

One possible role for sORF-encoded peptides that are similar to known proteins is to mimic the similar protein to interfere with its function. MiPs (or siPEPs) are important modulators of protein– protein and protein–DNA interactions that, for example, prevent the formation of functional protein complexes (Seo et al. 2013; Graeff et al. 2016). We suggest that the potential for sORFs that overlap with the CDS of protein-coding genes to be a source of small interfering peptides is currently underestimated (Fig. 6B). We found that approximately 30% of cis-SEPs harbor protein domains such as protein kinase domains and MYB-like DNA-binding domain or IDRs. The genes harboring CDS-sORFs were enriched in GO terms connected to protein binding and transferase activity. Also, some sORFs with disordered regions might mediate protein–protein or protein–nucleic acid interactions, as suggested previously (Mackowiak et al. 2015). Taken together, these findings suggest that sORFs may strongly interfere with protein interactions.

In this study, we explored several groups of sORFs, including those encoded by lncRNAs. The translation of peptides from lncRNAs is intriguing, and there is some evidence that these peptides play important biological roles in various processes (Kondo et al. 2010; Magny et al. 2013; Matsumoto et al. 2017). Nevertheless, the biological functions of most lncRNA-sORF-encoded peptides are currently unclear, especially those in the plant kingdom (Tavormina et al. 2015).

The transcription of the non-coding portions of the genome into lncRNAs is thought to give rise to the translation of sORFs located within them. In this case, some of these peptides would not be vital but may be important for survival under certain conditions by serving as a raw material for evolution (Fig. 6C). Knocking out select lncRNA-encoded peptides was not lethal in moss, but did influence moss growth under certain conditions. On the other hand, plants overexpressing an lncRNA-encoded peptide (41 aa) showed phenotypic differences compared to wild-type plants, suggesting a possible role for the lncRNA-encoded peptide in regulating cell growth and development. Our results lay the groundwork for systematic analysis of functional peptides encoded by sORFs.

The possible evolution of non-coding portions of the genome into protein-coding genes is also a subject of intensive debate (Carvunis et al. 2012; McLysaght and Guerzoni 2015; Couso and Patraquim 2017). According to our data, putative homologous sORFs tended to differ in length in most cases (Fig. 2D). Thus, we suggest that most sORFs expanded during evolution, providing support for the notion that they function as raw materials for selection; however, this point requires further confirmation.

## METHODS

### Physcomitrella patens growth conditions

*Physcomitrella patens* protonemata were grown on BCD medium supplemented with 5 mM ammonium tartrate (BCDAT) or 0.5% glucose during a 16-h photoperiod at 25°C in 9-cm Petri dishes (Nishiyama et al. 2000). For all analyses, the protonemata were collected every 5 days. The gametophores were grown on free-ammonium tartrate BCD medium under the same conditions, and 8-week-old gametophores were used for analysis. Protoplast was prepared from protonemata as described previously (Fesenko et al. 2015).

For morphological analysis, protonemal tissue 2 mm in diameter were inoculated on BCD and BCDAT 9-cm Petri dishes. For growth rate measurements, photographs were taken at 7 d intervals over 42 days. Protonemal tissues and cells were photographed using a Microscope Digital Eyepiece DCM-510 attached to a Stemi 305 stereomicroscope or Olympus CKX41.

### Identification of coding sORFs in the *P. patens* genome

To identify sORFs with high coding potential, the sORFfinder (Hanada et al. 2010) tool was utilized. Intron sequences and CDS were used as negative and positive sets, respectively. Additional details are described in the Supplemental Methods. To select for sORFs that are transcribed, located in the exons of transcripts, and have introns, a bed file was generated using a python script (GffParser.py) and intersected with exon positions extracted from a gff3 file of *P. patens* genome annotations. To identify intergenic-sORFs, the bed file was also intersected with transcribed regions determined based on our RNAseq data (Fesenko et al. 2017). Using an R script, sORFs fully overlapping with exons were selected; 75,685 sORFs remained after this step. Identical sORFs were removed from the dataset. In addition, sORFs overlapping repetitive regions identified by RepeatMasker, as well as sORFs comprising parts of annotated *P. patens* proteins, were also removed from the dataset, resulting in a final dataset of sORFs comprising 70,095 sequences.

### sORF classification

The step-by-step procedure performed for sORF classification is illustrated in Supplemental Fig. S14. In the first step, lncRNA-sORFs were identified by searching for identical sORFs in known lncRNA databases, including CantataDB (Szczesniak et al. 2016), GreenC (Paytuvi Gallart et al. 2016) and our previously published moss dataset (Fesenko et al. 2017). After this sORF bed file was intersected with moss genome annotation, the locations of the sORFs on transcripts were determined, resulting in the further classification of genic-sORFs into uORFs, dORFs, CDS-sORFs and interlaced-sORFs.

Because alternative splicing leads to inaccuracy in genome annotation, the locations of a subset of genic-sORFs cannot be unambiguously classified, as they can be located in different regions in different isoforms of the same gene. All sORFs located on transcripts that were not annotated in the *P. patens* genome but were identified using our RNAseq data were classified as intergenic-sORFs.

To detect alternatively spliced sORFs (AS-sORFs), a bed file with sORF locations was intersected with a bed file containing intron coordinates for all isoforms. Those sORFs that overlapped for both exons (see above) and introns were classified as AS-sORFs.

### Evolutionary conservation analysis

The transcriptomes of nine plant species were downloaded from Phytozome v12: *Sphagnum fallax* (release 0.5), *Marchantia polymorpha* (release 3.1), *Selaginella moellendorffii* (release 1.0), *Spirodela polyrhiza* (release 2), *Arabidopsis thaliana* (TAIR 10), *Zea mays* (Ensembl-18), *Oryza* sativa (release 7), *Volvox carteri* (release 2.1) and *Chlamydomonas reinhardtii* (release 5.5). The transcriptome of *Ceratodon purpureus* was *de novo* assembled using Trinity (Haas et al. (2013)). To identify transcribed homologous sequences, tBLASTn (word size = 3) was performed using sORF peptide sequences as queries and the transcriptome sequences of the abovementioned species as subjects. The following cutoffs parameters were used to distinguish reliable alignments: E-value < e-5 and query coverage > 60%. Our E-value cutoff was obtained by applying a multiple comparison correction (Bonferroni correction) of 0.05, which is commonly used in biological experiments.

Pairwise *K*_*A*_/*K*_*S*_ ratios were calculated using the codeml algorithm with PAML software (Yang 2007). The calculation procedure, which was facilitated using a custom-made python script (protein_Ka_Ks_codeml.py), included alignment extraction from the tBLASTn output, PAL2NAL (Suyama et al. 2006) correction of the nucleotide alignment using the corresponding aligned protein sequences and calculation of *K*_*A*_/*K*_*S*_ ratios using codeml. The script implements packages from biopython (Cock et al. 2009). To estimate homologous sORF lengths, a python script (sORF_completeness_v2.0.py) was designed. Additional details are described in the Supplemental Methods.

### GO enrichment analysis

GO enrichment analysis was performed using the topGO bioconductor R package using the Fisher’s exact test in conjunction with the ‘classic’ algorithm (false discovery rate [FDR] < 0.05). Gene Ontology (GO) terms assigned to *P. patens* genes were downloaded from Phytozome. Only GO terms containing >5 genes in a background dataset were considered in the enrichment analysis. Redundant GO terms were removed using the web-based tool REVIGO (Supek et al. 2011).

### Peptide and protein extraction

Endogenous peptide extraction was conducted as described previously (Fesenko et al. 2015). Proteins were extracted as described previously (Fesenko et al. 2016). Additional details are described in the Supplemental Methods.

### Mass-spectrometry analysis and peptide identification

Mass-spectrometry analysis was performed using three biological and three technical repeats for the proteomic (Fesenko et al. 2017) and peptidomic datasets. Analysis was performed on two different mass spectrometers: a TripleTOF 5600+ mass spectrometer with a NanoSpray III ion source (ABSciex, Canada) and a Q Exactive HF mass spectrometer (Q Exactive HF Hybrid Quadrupole-Orbitrap mass spectrometer, Thermo Fisher Scientific, USA). Additional details are described in the Supplemental Methods.

All datasets were searched individually with MaxQuant v1.5.8.3 (Tyanova et al. 2016) against a custom database containing 32926 proteins from annotated genes in the latest version of the moss genome (V3.1, (Lang et al. 2018)), 85 moss chloroplast proteins, 42 moss mitochondrial proteins and 72095 predicted sORF peptides. MaxQuant’s protein FDR filter was disabled, while 1% FDR was used to select high-confidence PSMs, and ambiguous peptides were filtered out. Moreover, any PSMs with Andromeda scores of less than 30 were discarded (to exclude poor MS/MS spectra). For the dataset of endogenous peptides (named “peptidomic”, Supplementary Table S2), the parameter “Digestion Mode” was set to “unspecific” and modifications were not permitted. All other parameters were left as default values. For the dataset of tryptic peptides (named “proteomic”) the parameter “Digestion Mode” was set to “specific” (the Trypsin/P), MaxQuant’s protein FDR filter was disabled, and the peptide FDR remained at 1 %. All other parameters were left as default values. Features of the PSMs (length, intensity, number of spectra, Andromeda score, intensity coverage and peak coverage) were extracted from MaxQuant’s msms.txt files.

To filter out MS peptides that do not provide unambiguous evidence of sORF peptide expression, we assessed the number of times a peptide occurred in the whole moss genome by searching for exact matches to the MS peptides in the six-frame translated genome. Of 629 unique peptides, 595 peptides (corresponding to 570 sORFs) matched only to the corresponding sORF peptide in the translated genome. The moss genome has a number of paralogous genes that resulted from two whole-genome duplication events (Lang et al. 2018). MS peptides from such paralogous sORFs will be discarded if they match to more than one locus in the genome. To prevent this, we identified paralogous sORFs in the moss genome by tBLASTn and aligned their coordinates with the multi-hit MS peptide coordinates. This identified 15 MS peptides (14 sORFs) that matched to paralogous sequences and were discarded from further analysis. Our final high-confidence set included 584 translatable sORFs.

### RT-PCR analysis of AS-sORFs

Total RNA from gametophores, protonema and protoplasts was isolated as previously described (Cove et al. 2009). RNA quality and quantity were evaluated via electrophoresis in an agarose gel with ethidium bromide staining. The precise concentration of total RNA in each sample was measured using a Quant-iT^TM^ RNA Assay Kit, 5–100 ng on a Qubit 3.0 (Invitrogen, US) fluorometer. The cDNA for RT-PCR was synthesized using an MMLV RT Kit (Evrogen, Russia) according to the manufacturer’s recommendations employing oligo(dT)17-primers from 2 µg total RNA after DNase treatment. The primers were designed using Primer-BLAST (Ye et al. 2012) (Supplementary Table). The minus reverse transcriptase control (-RT) contained RNA without reverse transcriptase treatment to confirm the absence of DNA in the samples. The RT-PCR products were resolved on an 1.5% agarose gel and visualized using ethidium bromide staining.

### Generation of overexpression and knockout lines

To obtain overexpression lines of the PSEP1 (Pp3c9_sORF1544), PCR was carried out using genomic DNA as a template and PEP4f and PEP4r primers (Supplemental Table S6). Amplicons were cloned into the pPLV27 vector (GenBank JF909480) using the Ligation-independent (LIC) procedure (Aslanidis and de Jong 1990; De Rybel et al. 2011). The resulting plasmid was named pPLV-Hpa-4FR and used for transformation. Additional details are described in the Supplemental Methods.

PSEP1 (sORF Pp3c9_sORF1544), PSEP3 (Pp3c25_sORF1253) and PSEP25 (Pp3c25_sORF1000) knockout lines were created using the CRISPR-Cas9 system (Collonnier et al. 2017). The coding sequences were used to search for CRISPR RNA (crRNA) preceded by a *S. pyogenes* Cas9 PAM motif (NGG) using the web tool CRISPR DESIGN (http://crispr.mit.edu/). The crRNA closest to the translation start site (ATG) was selected for cloning (Supplemental Table S6).

Protoplasts were transformed using PEG transformation protocol (Schaefer and Zryd 1997). Additional details are described in the Supplemental Methods. The plasmids pACT-CAS9 (for CAS9 expression) and pBNRF (resistance to G418) were kindly provided by Dr. Fabien Nogué.

Independent knockout and overexpression mutant lines have been obtained (Supplemental Figs. S10-12).

The ploidy level of the *PSEP1* overexpression and *psep1* knock-out lines were estimated using flow cytometry. Protoplasts were fixed in cold 70 % methanol, washed in TBS with 0.1 % Triton X-100, then washed with TBS and stained with 500 ng/ml DAPI. The fluorescence was analyzed with a flow cytometer NovoCyte (ACEA Biosciences) and Novoexpress data software. Fluorescence was excited at 405 nm, and detection was at 445/45 nm.

## DATA ACCESS

All raw mass spectrometry data from this study have been deposited to the ProteomeXchange Consortium via the PRIDE (Vizcaino et al. 2016) partner repository with the dataset identifiers PXD005223, PXD007922, PXD007923, PXD007973.

## SOFTWARE AVAILABILITY

All data were analyzed using Python (http://www.python.org, v 3.5), and R (http://www.R-project.org, R Development Core Team 2006). All scripts are available at Zenodo (doi: 10.5281/zenodo.1160331) and are maintained in the GitHub code repository: https://github.com/Kirovez/Scripts_sORFs_MS.

## ACKNOWLEDGEMENTS

This work was supported by the Russian Science Foundation (project No.17-14-01189). Some of mass spectrometric measurements were performed using the equipment of the “Human Proteome” Core Facility of the Orekhovich Institute of Biomedical Chemistry (Russia) which is supported by the Ministry of Education and Science of the Russian Federation.

## Authors’ contributions

IF and IK conceived and designed experiments. AK performed moss transformation experiments. IF, RK, VL, DK, EG, VZ, IB and AM performed the proteomics analyses. IF, IK and GA performed the statistical and bioinformatics analyses. IF, IK, VI and VG wrote the manuscript with input from all authors. IF supervised the project. All authors read and approved the final manuscript.

